# Mechanisms Controlling Hemolymph Circulation Under Resting Conditions In The Chagas Disease Vector *Rhodnius Prolixus* (Stål)

**DOI:** 10.1101/2023.10.19.563123

**Authors:** María José Villalobos Sambucaro, María Eugenia Alzugaray, Jorge Rafael Ronderos

## Abstract

Chagas disease vectors can ingest several times its own volume in blood with each meal. This ad libitum feeding causes an intense process of diuresis inducing the insect to eliminate a large quantity of urine during the next few hours. This process, which is under the control of endocrine and neuroendocrine systems is necessary to restore homeostasis, and to begin physiological mechanisms leading to growth and reproduction. To ensure diuresis, the speed of circulation of the hemolymph must be increased to allow the Malpighian tubules to produce the urine. Behind this acute phenomenon, triatominae insects can spend several weeks without feeding. In this way, it could be assumed that during most of the time of life the insect is in a resting condition. Triatominae circulatory system is quite simple, including a dorsal vessel which pumps hemolymph in an anterograde direction. The return is caused by peristaltic contractions of the anterior midgut. While the mechanisms controlling the circulation of the hemolymph during post-prandial diuresis was largely analysed, the mechanisms controlling it during resting conditions is poorly understood. In this study we analyse several canonical pathways (i.e. L-type VGCC; GPCR; RyR; IP3R) and a novel system represented by the recently characterized Piezo proteins. Our results show that during the resting condition hemolymph circulation depends on a cross-talk between myogenic activity, inhibitory and stimulatory cell messengers, and also Piezo proteins. We present for the first time the existence of a putative Piezo protein in Hemiptera.

## INTRODUCTION

Triatominae insects ingest in every meal many times its own volume in blood. In fact, it was shown that *Rhodnius prolixus* first instar larvae can ingests up to 12 times its weight in a few minutes (Buxton, 1930). Consequently, the insect is challenged with sudden changes, such as an increase on its weight, a misbalance of ionic equilibrium, an excess of water, changes of the osmotic concentration, etc. Due that, the insect eliminates a high volume of urine during more than 180 min (Maddrell, 1963, 1964a). This process is drived by the activity of diuretic hormones, as serotonin (5-HT), which turn on Malpighian tubules (MTs) (Maddrell et al., 1991, 1993a). In fact, 5-HT is released from the mesothorax ganglion to the hemolymph immediately after the insect begins to feed (Maddrell, 1964b). The urine produced by MTs drains into the rectum together with faeces coming from the posterior midgut, where they are mixed by peristaltic movements and periodically eliminated (Santini and Ronderos, 2007). This process constitutes the main natural way of Chagas disease transmission (https://www.paho.org/en/topics/chagas-disease).

It was shown in the related species *Triatoma infestans*, that MTs secrete the neuropeptide Allatotropin (AT) which stimulates muscle wall of the rectum inducing peristaltic contractions and voiding (Santini and Ronderos, 2007, 2009a, 2009b). MTs are no innervated, being modulated in a neuroendocrine way. Consequently, to ensure the diuretic activity, the secretion and distribution of hormones must be accompanied by the increase of the speed of hemolymph recirculation. To ensure that, a number of powerful peristaltic movements of the anterior midgut (crop) are the main factor causing a fast movement of the hemolymph in an antero-posterior direction (Maddrell, 1964a). The recirculation is complemented by an increase of the frequency of contractions of the dorsal vessel (DV) (Sterkel et al., 2010; Villalobos Sambucaro et al., 2015). Regarding this, we have shown that, during the physiologically critical period of diuresis, at least three different cell messengers are involved in a highly synchronized cross-talk. We have shown in *T. infestans*, as well as, in *R. prolixus* that serotonin triggers an increase of both aorta contractions (driving hemolymph in an anterograde way), and the frequency of peristaltic contractions of the crop (to move hemolymph in the opposite direction). This process is synergized by AT (Sterkel et al., 2010; Villalobos Sambucaro et al., 2015). It was also shown in both, *T. infestans* and *R. prolixus*, that the crop, as well as the cephalic portion of the aorta are innervated with allatotropic fibres (Masood and Orchard, 2014; Riccillo and Ronderos, 2010; Sterkel et al, 2010). Furthermore, the presence of allatotropic open-type cells in the epithelial sheet of the anterior region of the crop of *T. infestans* were proved, suggesting the existence of a paracrine mechanism modulating peristaltic activity (Riccillo and Ronderos, 2010; Sterkel et al., 2010). This process is also influenced by Allatostatin-C (AST-C) (Kramer et al., 1991), which counteracts the stimulus of AT decreasing both, the rate of peristaltic waves of the crop and the frequency of contractions of the aorta diminishing the availability of cytosolic calcium (Villalobos et al., 2016). It was also shown that both organs express receptors for all these three messengers (i.e. 5-HT, AT and AST-C) (Paluzzi et al., 2015; Villalobos Sambucaro et al., 2015, 2016).

As we stated above, hemolymph is moved in a postero-anterior direction by mean of the dorsal vessel (DV) contractions. The DV are constituted by the heart, located at the final segments of the abdomen, associated to the alary muscles on lateral sides (Fig. 1 A-C); and the aorta which runs throughout the abdomen up to the thorax, opening at the level of the head (Fig. 1 C-D) (Chiang et al., 1990; Villalobos Sambucaro et al., 2015). The heart and the aorta present little holes (ostia) by which hemolymph enter to the circuit from the hemocele to be pumped (Fig. 1 A and D). Unlike some species, in which the flow direction through the DV alternates between postero-anterior and antero-posterior (Gerould, 1933; Glenn et al., 2010; League et al., 2015) in triatominae the DV normally pumps only in anterograde direction (Villalobos Sambucaro et al., 2015).

**Figure 1.**
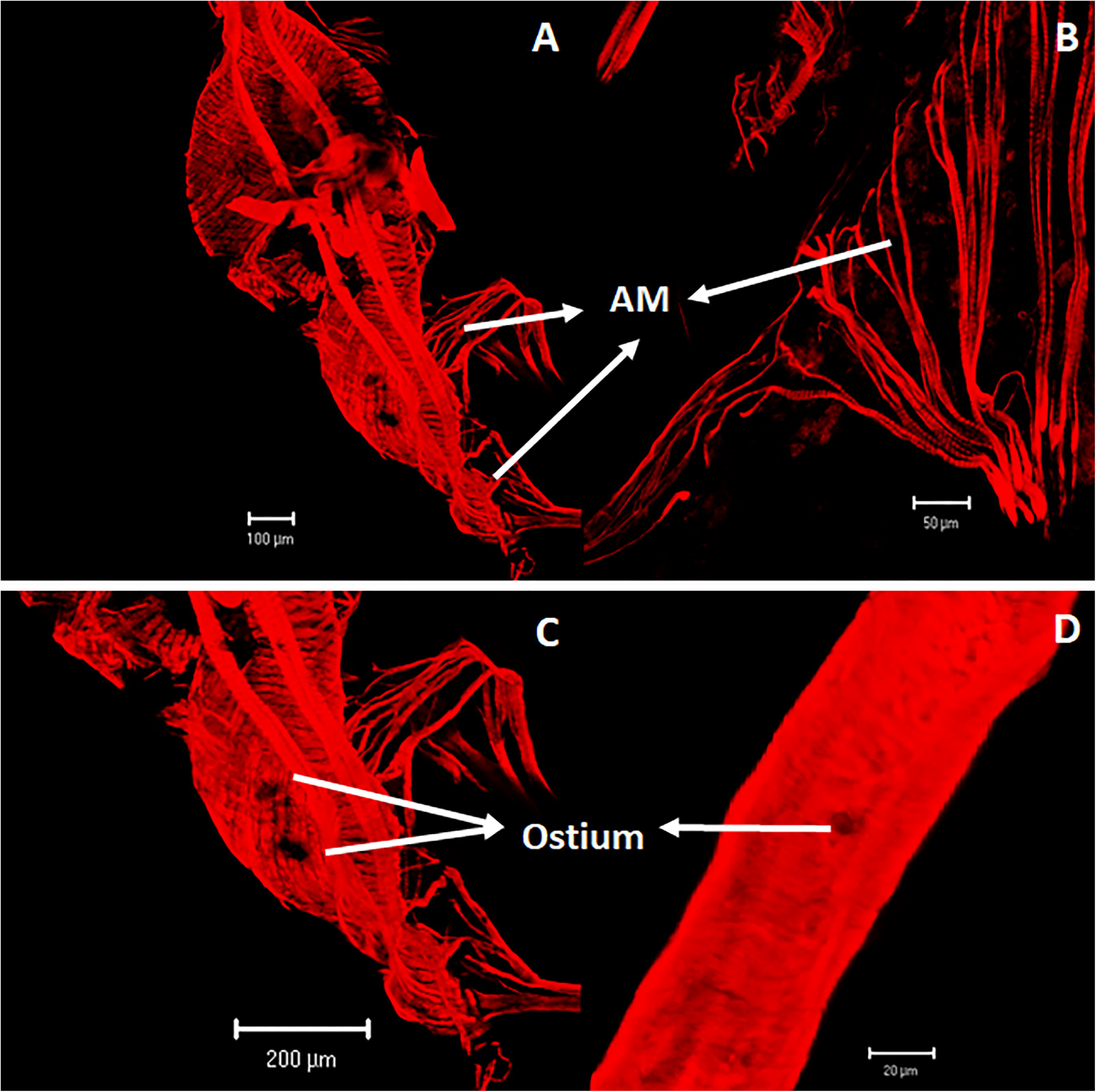
Dorsal vessel and associated alary muscles of an adult male of *R. prolixus* labelled with rhodamine-phalloidin to visualize muscle fibres organization. A) Heart and alary muscles. B) Detailed view of an alary muscle. C) and D) Detailed views of the heart and aorta showing the ostia through which the hemolymph goes into the circulatory system to be pumped.

Triatominae insects can spend long times without feed so, post-prandial diuresis is an acute critical process, occurring few times along the life. In the present study, we attempted an approach to understand how cardiac and crop muscle activity is controlled during resting condition, which represents the longest period during the life of the insect. We analysed different canonical ways that might be acting to maintain a beating rhythm to sustain hemolymph circulation during periods in between consecutive meals. We show that, beside the existence of myogenic activity, the rate of contractions is influenced by messengers of peptidic nature such as Allatotropin and Allatostatin-C, but also by proteins susceptible to mechanical forces associated to plasma membranes that modulates electric activity of the cell; the piezo-proteins.

Piezo proteins, a family of plasma membrane associated proteins, have been identified few years ago. This family of proteins, originally characterized in mice (Piezo1 and Piezo2), act as cation channels (Coste et al., 2010). They are also present in *Drosophila melanogaster* (Kim et al., 2012). Indeed, in *D. melanogaster*, it has been shown that piezo-like proteins would modulate intracellular Ca^2+^ concentration in the heart (Zechini et al., 2022). Based on experimental designs involving Piezo protein agonists (Jedi 1 and Jedi 2) we found that both, DV contractions, as well as the peristaltic movements of the crop depend also on piezo-like proteins. Moreover, we found that *R. prolixus* genome predicts the expression of a protein that presents homology with other piezo proteins previously characterized.

Mechanism controlling hemolymph circulation under resting conditions involves a complex system of signals involving myogenic activity, myoregulatory peptides, and reflex contractions induced by piezo proteins.

## MATERIAL AND METHODS

### Insects

Adult males of *R. prolixus* were obtained from a colony maintained at 28 ± 2°C, 30% relative humidity, and 12:12 hour light-dark period. Males were identified during 5^th^ instar larva, isolated and starved during 10 days before meal was offer. Only those individuals fed *ad libitum* were selected for experiments. After moult, adult males were starved for at least 21 days to ensure that the crop was empty.

### Muscle tissue labelling with phalloidin

Dorsal vessel was dissected under binocular microscope and fixed in formaldehyde-phosphate buffered saline (PBS) (4%) at 4°C for 12 hours. Samples were washed in PBS-Tween (0.05%) and then permeabilized with Triton X-100 (1%) (24 hours at 4°C). To visualize the arrangement of the muscle tissue, samples were then incubated overnight at 4°C with a rhodamine-phalloidin solution (Sigma–Aldrich) (1/1000) and finally mounted in Vectashield. Samples were analysed with a confocal laser scan microscope (LSM) Zeiss LSM 510 Meta (Adami et al., 2011, 2012a, 2012b; Alzugaray et al., 2013).

### Myoregulatory assays

Assays were designed to study the frequency of contractions of the DV and the rate of peristaltic contractions of the crop *in vivo* as in previous studies (Sterkel et al, 2010; Villalobos Sambucaro et al, 2015, 2016). After emergence insects were starved during 20 to 25 days after moulting, stablishing as resting condition the moment when the crop was empty, to ensure that physiological processes associated to digestion were finished.

Briefly, each individual was placed under a stereoscopic microscope with a digital camera annexed. The legs were hold with plasticine on a Petri dish and the wings removed, to allow aorta and crop visualization through the translucent dorsal cuticula. All the treatments were diluted in *R. prolixus* saline (modify from Maddrell 1993) and applied directly into the abdomen through a little incision in the conexive using a Hamilton micro-syringe (5µl). To minimize the effect of the stress caused by handling, insects were allowed to rest for 30 minutes before the treatment administration. The contractions of the aorta and crop were observed through the dorsal cuticle (segments III to V of the abdomen). The number of contractions in a 3-min period was recorded at 5, 15 and 30 minutes after treatments, all data were recorded by the same operator. Results are expressed as number of contractions/min.

### Calcium involvement on dorsal vessel and crop frequency of contractions

Different compounds previously assayed in our laboratory have been used to study the relevance of Ca^2+^ on the activity of the aorta and crop (Alzugaray and Ronderos 2018; Alzugaray et al., 2019, 2021). The drugs employed and the concentration used are listed below:

- EDTA (Sigma-Aldrich) -40 mM-. A calcium chelator that does not penetrate the cell.
- BAPTA/AM. 1, 2-bis (2-aminophenoxy) ethane-N, N, N‘, N‘-tetra-acetic acid (Tocris Bioscience) -300 µM and 400 µM-. A high-affinity and membrane permeable (intracellular) Ca^2+^ chelator (Tsien, 1981).
- Nifedipine (Sigma-Aldrich) -30µM and 60µM-, is a blocker of L-type voltage sensitive calcium channels (Guazzi et al., 1977; Zsotér and Church, 1983).
- Xestospongin-C (Xe-C) -6 µM-(Tocris Bioscience). A membrane-permeably compound isolated from the sponge *Xestospongia sp*., that inhibit Inositol 1,4,5 triphosphate calcium receptor/channel (IP_3_R) in the endoplasmic reticulum (ER) (Gafni et al., 1997; Kiselyov et al., 1998).
- Ryanodine (Ry) -10 μM; 10 nM-(Tocris Bioscience). A diterpenoid alkaloid of plant origin that specifically modifies the activity of the Ryanodine receptor calcium channels (RyR) in the ER, maintaining them in an open state at nanomolar concentrations, and closed at micromolar concentrations (Bull et al., 1989; Meissner, 1986).

### Aorta and crop contractile activity associated with GPCRs transductional pathways

In order to test if GPCRs signalling cascade could be modulating the basal frequency of the organs under study, we assayed the effect of the following drugs:

- Sulfhydrylreactive hydrobromide (SCH-202676) (Tocris Bioscience) -1, 5, 10µM-. SCH-202676 reversibly inhibits agonist and antagonist binding to a variety of GPCRs in mammalian and non-mammalian cells (Alzugaray et al., 2021, Fawzi et al., 2001; Gao et al., 2004; Hartz et al., 2008; Lewandowicz et al., 2006).
- U73122, an inhibitor of phospholipase-C (PLC) (Tocris Bioscience) -10µM-. PLC is an enzyme that produces IP_3_ when is activated through a GPCR coupled to a G_αq_, resulting in an increase of cytosolic Ca^2+^ level by activation of IP_3_R in the ER (Bleasdale et al., 1990; Smith et al., 1990).

### Allatotropin and Allatostatin-C interaction as myoregulators during resting condition

Looking for the probable interaction between AT (Kataoka et al., 1989), which induces an increase of aorta contractions (Sterkel et al., 2010; Villalobos Sambucaro et al., 2015) and AST-C that antagonises its activity (Villalobos Sambucaro et al., 2016) we assayed the effect of AT (10^-9^ M) (Center for Biotechnology Research, Kansas State University) (Li et al., 2003), in the presence of an AST-C-antiserum (1/500) (Genemed Synthesis (San Francisco, Calif., USA)) (Li et al., 2004).

### Relevance of Piezo proteins and reflex contractions on hemolymph circulation during resting

In order to analyse the involvement of mechanical factors and Piezo proteins in the myoregulatory control of aorta and crop, we assayed the effect of two agonists:

- Jedi 1 and Jedi 2 (Sigma-Aldrich) -250 μM and 2.5 sM respectively-. Piezo1 chemical activators (Wang et al., 2018).

### Identification and bioinformatic characterization of the Piezo putative channels

The sequences corresponding to mouse Piezo 1 (ADN28064) and Piezo 2 (ADN28065) Mus musculus (Chordata: Mammalia), were used to look for putative piezo proteins in the genome of *R. prolixus* (https://www.vectorbase.org), and several phyla of Opisthokonta (https://blast.ncbi.nlm.nih.gov/Blast.cgi). The sequences corresponding to Piezo 1 and Piezo 2, the species included and their accession numbers are: *R. prolixus* (Arthropoda) RPRC014024; *Lingula anatina* (Brachiopoda) XP_013404042; *Owenia fusiformis* (Annelida) CAH1780572; *Octopus sinensis* (Mollusca) XP_029647893; *Trichoplax sp*. (Placozoa) RDD46920; *Hydra vulgaris* (Cnidaria) XP_047142278; *Rozella allomycis* (Fungi) EPZ31590; *Amphimedon queenslandica* (Porifera) XP_019849310; *Beroe abyssicola* (Ctenophora) AQX17753; *Salpingoeca rosetta* (Choanoflagellata) XP_004997271.

Looking for Piezo protein characteristic domains, the sequence was further analysed with InterproScan (www.ebi.ac.uk/interpro/result/InterProScan/#table). The transmembrane domains analysis were performed by using TMHMM – 2.0 (https://services.healthtech.dtu.dk/services/TMHMM-2.0/).

The selected sequences were aligned using Clustal Omega multiple sequence alignment program (https://www.ebi.ac.uk/Tools/msa/clustalo/) and post analysed with JalView 2.7 (Waterhouse et al., 2009).

Finally, the probable phylogenetic relationships and the evolutionary analysis were performed by mean of the Maximum Likelihood methodology (500 bootstrap replicates) by using MEGAX64 software (Tamura et al., 2013).

### Statistical analysis

Significant differences were evaluated by One Way Analysis of Variance (ANOVA). Single post-hoc comparisons were tested by the LSD test. Only differences equal or less than 0.05 were considered significant. All experimental groups were constituted by 5 to 10 individuals. Data are expressed as means ± standard error.

## RESULTS

### Relevance of extracellular and cytosolic calcium on the contractile activity of the aorta

As a first approach to test the relevance of cytosolic calcium on the frequency of contractions of the aorta we analysed the effect of BAPTA/AM, a permeable chelator, in two different concentrations. The results show that at both concentrations (i.e. 300 and 400 µM) the frequency of contractions was diminished (being the effect longer with the highest concentration assayed) showing that an increase of cytosolic calcium is necessary to cause the contraction of the aorta (Fig. 2A and B). The use of an extracellular chelator as EDTA, completely avoid aorta contraction suggesting that a Ca^2+^ influx from the extracellular matrix is necessary to cause a contraction (Fig. 2C).

**Figure 2.**
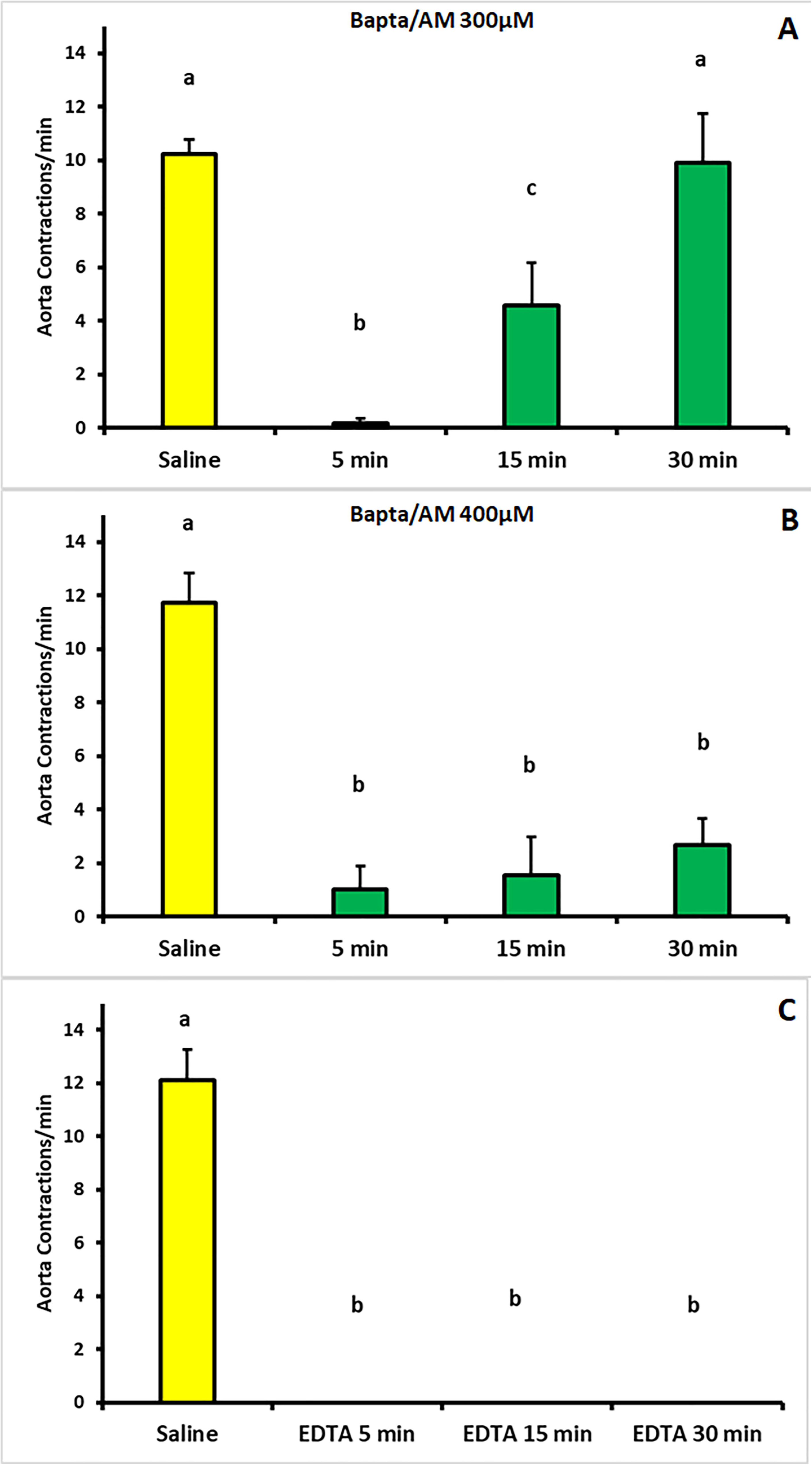
Aorta response to chelators. A) and B) The use of two different concentrations of BAPTA/AM (a permeable chelator actin in the cytosol) show that an increase of Ca^2+^ concentration in the cytosol is necessary for aorta contraction. C) The use of an external chelator as EDTA suggests that an extracellular influx of calcium is necessary for the contractile activity of the aorta. Bars represent mean ± SE. Different letters mean statistically significant differences.

### Extracellular calcium influx through L-Type VGCC and the opening of Ryanodine receptors in the endoplasmic reticulum

In view of the results obtained by the use of EDTA, showing the relevance of the extracellular source of Ca^2+^, we decide to check one of the most common pathways of calcium entry. To do that, we assayed the compound Nifedipine; a molecule that selective blocks L-Type voltage-gated calcium channels (VGCC) located at the plasma membrane. Results showed a statistically significant transient decrease of the frequency of contractions, confirming the importance of the extracellular Ca^2+^ influx. Indeed, these results suggest that L-Type VGCC are a relevant pathway during muscle contraction of the aorta (Fig. 3A).

**Figure 3.**
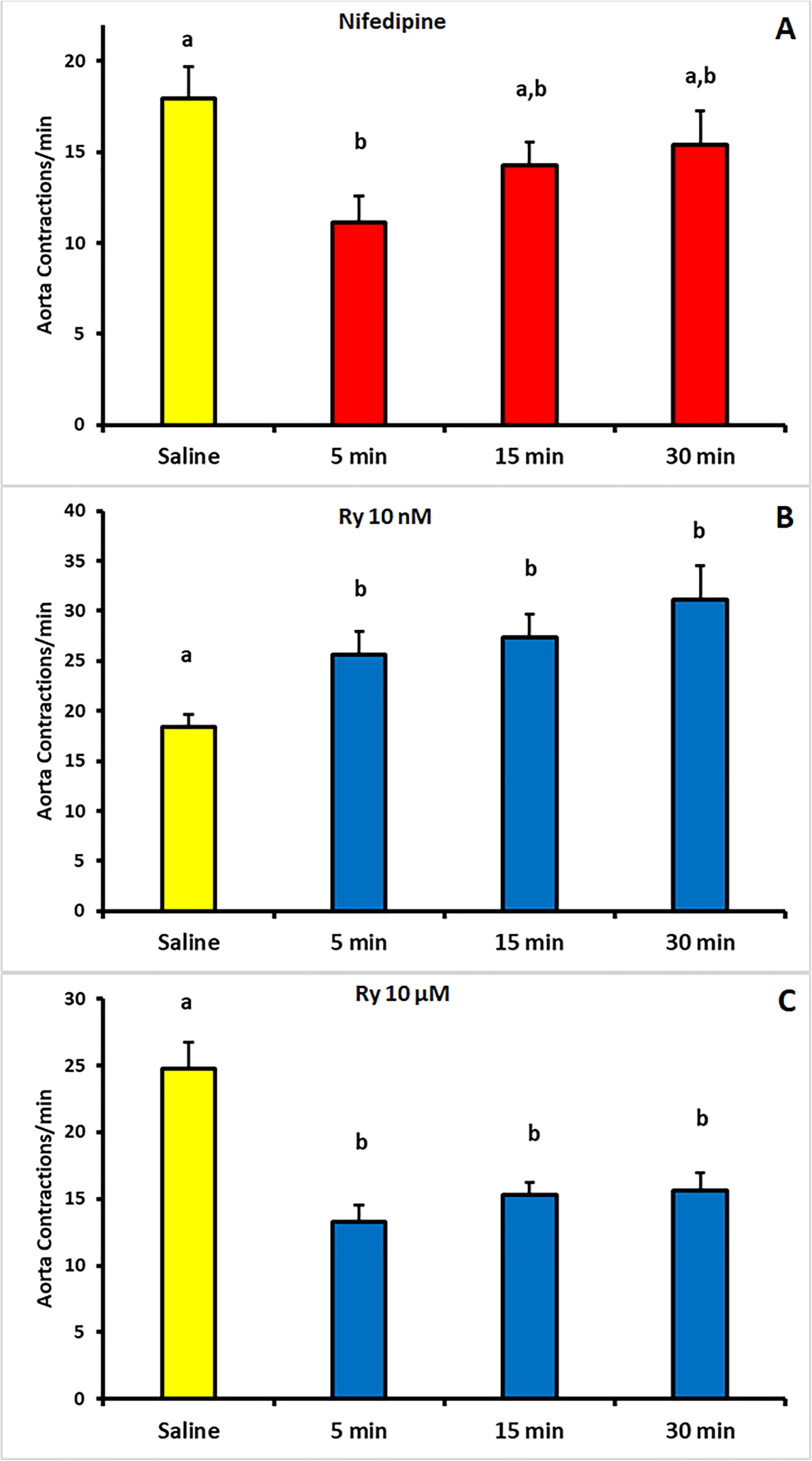
Calcium influx through L-type voltage gate calcium channel. A) Nifedipine, a specific VGCC blocker transiently inhibits the frequency of contractions. B) and C) The use of Ryanodine at inhibitory concentrations also decreases the contractile activity of the aorta suggesting that action potentials are involved. Each bar represents mean ± SE. Different letters mean statistically significant differences.

Calcium entry induces an increase of the cytosolic concentration of the ion. A common associated response is the opening of Ryanodine receptors (RyRs) channels at the level of the smooth endoplasmic reticulum (ER). RyRs opening is modulated by Ca^2+^. RyRs specifically respond to the vegetal diterpenoid Ryanodine (Ry), being stimulated at nanomolar concentrations (facilitating the outcome of Ca^2+^ from the SR) and inhibited when treated with micromolar concentrations. Our results showed that Ry 10 nM decreased the contractile activity, suggesting that Ca^2+^ influx through VGCC facilitates the increase of cytosolic calcium through the opening of RyR to induce muscle contraction (Fig. 3C). As a control of the reaction we performed a similar assay challenging the aorta to a stimulatory concentration of Ry (10 μM) (Fig. 3B).

### GPCRs and the IP_3_/IP_3_R pathway

Due that both treatments (i.e. Nifedipine and Ry) produce a decrease but do not completely avoid the contractile activity of the aorta, we decided to check the existence of a response mediated by extracellular messengers. Taking into account that the activity of numerous cell messengers is mediated by GPCRs, which involve the IP_3_/IP_3_R pathway to increase the cytosolic Ca^2+^ concentration, we assayed the effect of Xe-C. This compound originally isolated from the sponge *Xestospongia exigua*, selectively inhibits the opening of IP_3_R at the level of ER. Again, a statistically significant decrease of the frequency of contractions was observed (Fig. 4A). Interestingly, the use of U73122, a compound that inhibits the activity of PLC which is involved in the synthesis of IP_3_, did not cause any effect (Fig. 4B). To check the proper activity of the compound, we performed a similar experiment using fed individuals. In this case the activity of the aorta was diminished, proving that U73122 effectively alters PLC activity in this species (data not shown).

**Figure 4.**
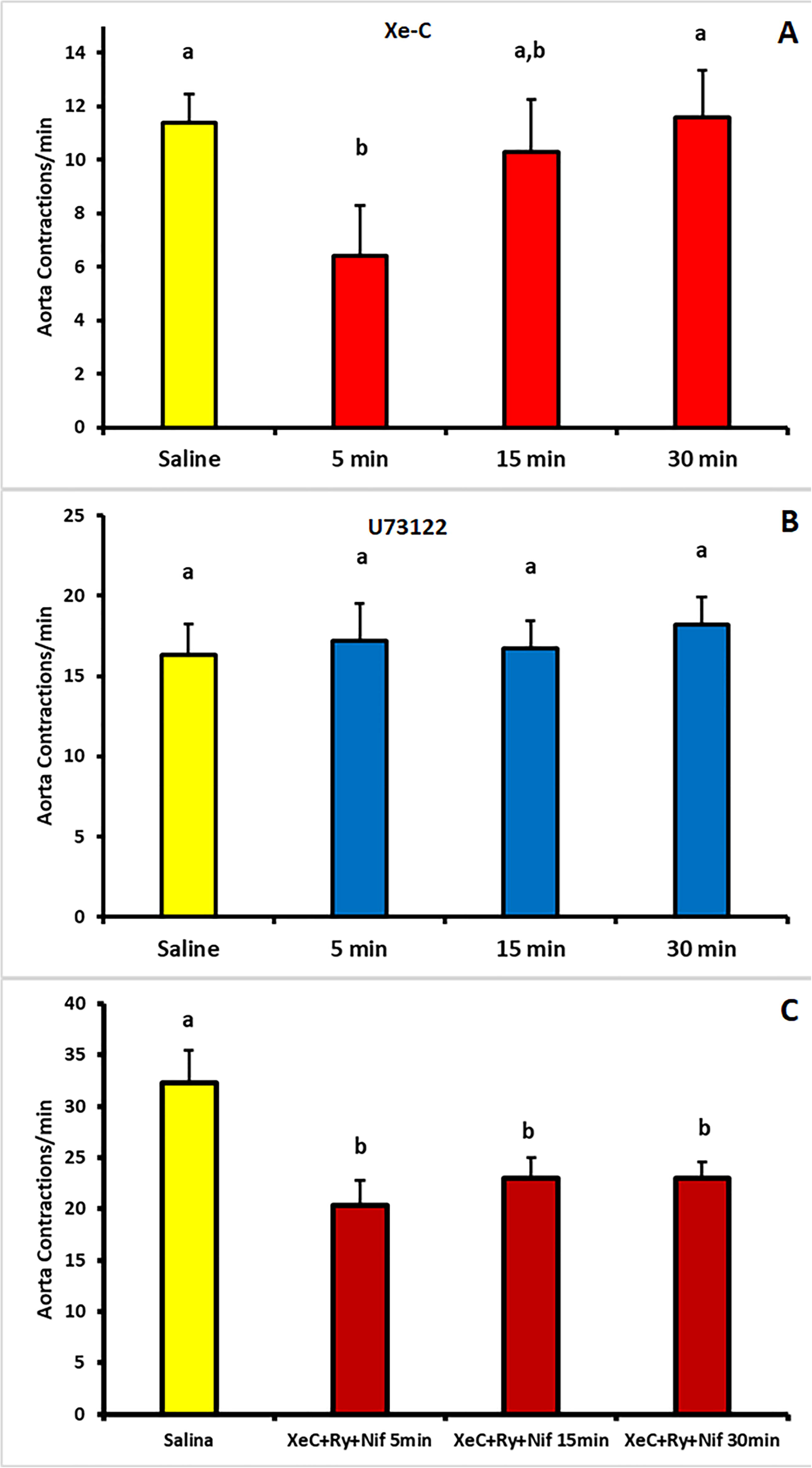
GPCR signalling involved on aorta contraction. A) Xestospongine-C inhibits IP_3_ dependent Ca^2+^ release from the endoplasmic reticulum suggesting that a signal related to GPCRs are involved. B) The use of a compound that blockades the activity of PLC does not alter the contractile activity of the aorta. C) Individuals treated with compounds that alter both L-type VGCC/RyR and IP_3_/IP_3_R simultaneously do not completely avoid aorta contractions. Each bar represents mean ± SE. Different letters mean statistically significant differences.

As it was shown, neither the compounds altering pathways of RYRs mediated by L-type VGCC nor those one acting on GPCR/IP_3_R pathways, completely blockade aorta frequency of contractions. This facts suggest that both of them could act in a complementary fashion regulating the activity of the dorsal vessel. To attempt a probable explanation, we decide to perform a new experiment evaluating the blockade of both pathways. To do that, we assayed the effect of Xe-C/Nifedipine/Ry, trying to generate a complete inhibition of the system. Again, a decrease of the frequency of contractions of the DV was found but it was not completely blockaded (Fig. 4C), suggesting the presence of another pathway acting to ensure a minimal constant activity of the DV.

Taking into account that IP3R pathway seems to participate in the regulation of the activity of the aorta during resting conditions, suggesting the involving of cell messengers acting via GPCRs, we performed a new experiment using an allosteric modulator of binding to GPCR(Fawzi, et al., 2001). If a stimulatory signal is acting through IP_3_R pathway, the expected result would be the inhibition of the signal, diminishing the rate of aorta contractions. On the contrary, the use of SCH-202676 10 µM caused a statistically significant increase of the frequency suggesting the participation of an inhibitory cell messenger also acting through a GPCR (Fig. 5A).

**Figure 5.**
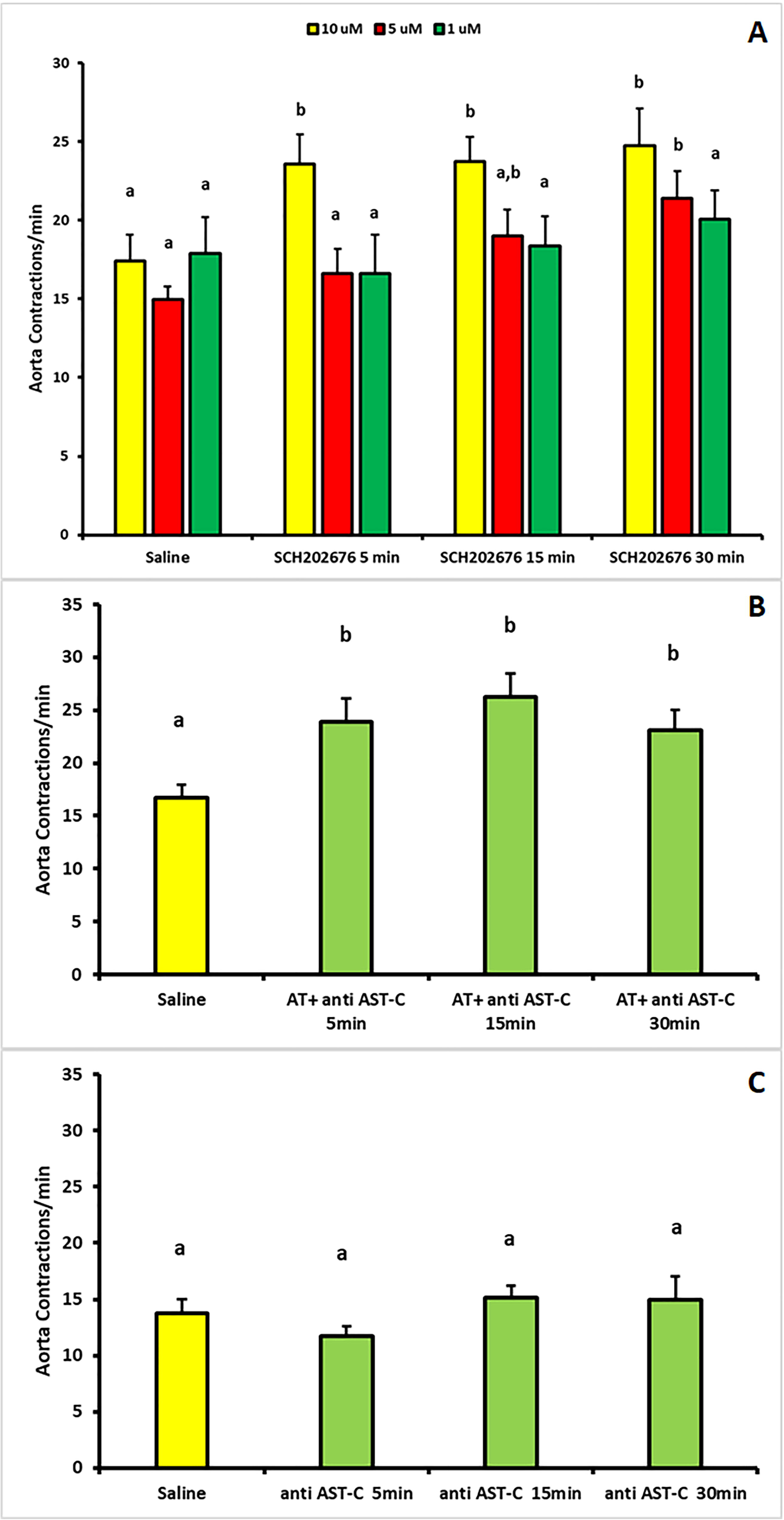
Allatotropin/Allatostatin-C activity also modulate the frequency of contractions of the aorta. A) The use of an allosteric modulator of GPCRs induces an increase of the number of contractions/min. B) In the presence of an AST-C-antiserum AT stimulates muscle contraction. C) Treatment of the individuals with the AST-C-antiserum does not modify the activity of the aorta. Bars represent mean ± SE. Different letters mean statistically significant differences.

### Myoregulatory peptides as modulators on the aorta frequency of contractions

Taking into account the results described above, we looked for the interaction of two independent signals, acting complementarily. Indeed, it was shown in *R. prolixus* that during post-prandial diuresis at least three messengers interact to regulate aorta frequency of contractions (serotonin, Allatotropin, Allatostatin-C). As it was previously proved, in the presence of 5-HT, AT increases the frequency of contractions, being this mechanism antagonized by AST-C (Villalobos Sambucaro et al., 2016). In this way, it might be possible that AT and AST-C interact to sustain the rhythm of contraction in a modulated fashion also during resting conditions. To check this hypothesis, we treated a group of unfed individuals with AT in the presence of an AST-C antiserum. A second group was treated only with the AST-C antiserum. The results showed that in the presence of AST-C antiserum, AT caused a statistically significant increase similar to the one observed in others physiological conditions (Villalobos Sambucaro et al., 2016, 2023) showing that AST-C is a canonical mechanism controlling the activity of AT (Fig. 5B). The group treated only with the AST-C antiserum did not show any difference with control group (i.e. saline injected individuals) (Fig. 5C).

### Peristaltic movements of the crop and the antero-posterior movement of the hemolymph

Regarding the hemolymph retrograde movement, it is caused by the peristaltic contractions of the crop (Maddrell, 1964a). As in previous studies, we also recorded the peristaltic rate of this organ which, receives and processes the blood during feeding. The results showed that the rate of contractions was only altered in insects treated with chelators (BAPTA/AM and EDTA) (Fig. 6A and B) and Ry (Fig. 6C). None of the other compounds assayed proved to be active; neither compounds acting on L-type VGCC, nor compounds that modulate GPCR response. These results suggest that a different mechanism is controlling peristaltic activity of the crop during resting conditions.

**Figure 6.**
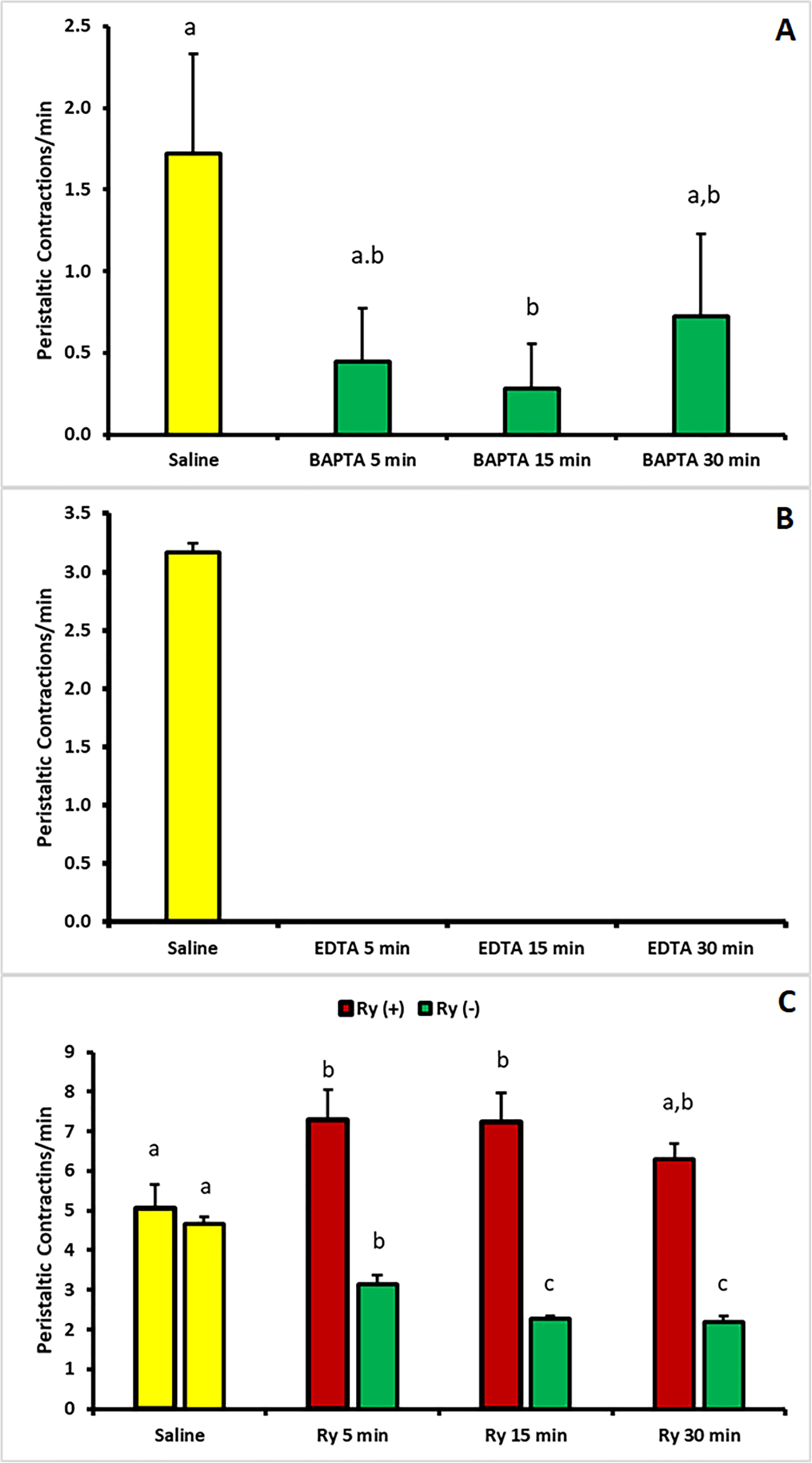
Peristaltic contractions of the anterior midgut (Crop) under resting conditions. A) The rate of peristaltic contractions is altered when cytosolic Ca^2+^ is chelated. B) Extracellular deplete of calcium completely avoid contractile activity of the crop. C) The use of Ryanodine also blockades the peristaltic activity of the crop. Bars represent mean ± SE. Different letters mean statistically significant differences.

### Piezo proteins in *R. prolixus* and its involvement in hemolymph circulation

A complex and complementary system involving both myogenic and peptidergic mechanisms seems to act to maintain the activity of the aorta during the resting conditions. However, as we stated above, neither GPCRs nor L-type VGCC related pathways seem to represent the complete machinery of the system modulating DV and crop activity. In fact, treatments to alter both pathways applied together, did not completely avoid the contractile activity of the dorsal vessel. Furthermore, the activity of the crop does not show rhythmic contractions. On the contrary, the low and variable peristaltic activity appears to be caused as a reflex response. To look further in the analysis of the complete machinery acting to modulate aorta and crop contractions during resting conditions, we analyse the probable involvement of piezo proteins.

An in-silico search for piezo-like proteins in *R. prolixus* showed that the genome predicts the expression of an orthologue (RPRC014024) presenting two domains considered characteristic of Piezo proteins. One of them from residue 1163 to 1294 (Coste et al., 2010). The second one, which in fact resemble a piezo-type mechanosensitive ion channel homolog (residues 1779 to 2368) (Coste et al., 2012) (Fig. 7A and B). The transcript, presenting 36 transmembrane domains show also a high degree of conservation with proteins of the family corresponding to several eukaryota groups, including non-metazoan organisms as Fungy and Choanoflagellata (Fig. 8 A and B). Indeed, the predicted protein includes the PFEW motif that it was proposed as a signature of this family of proteins (Prole and Taylor, 2013) (Fig. 7A). To delve into the evolutionary analysis, we looked for the degree of conservation of some other motifs (Fig. 7A and 8A). We found that the motif FLYRSPET, included in the probable piezo-type mechanosensitive ion channel at the C-terminal domain, is highly conserved in Bilateria, Cnidaria and Placozoa, and might be considered as a signature of this family of proteins.

**Figure 7.**
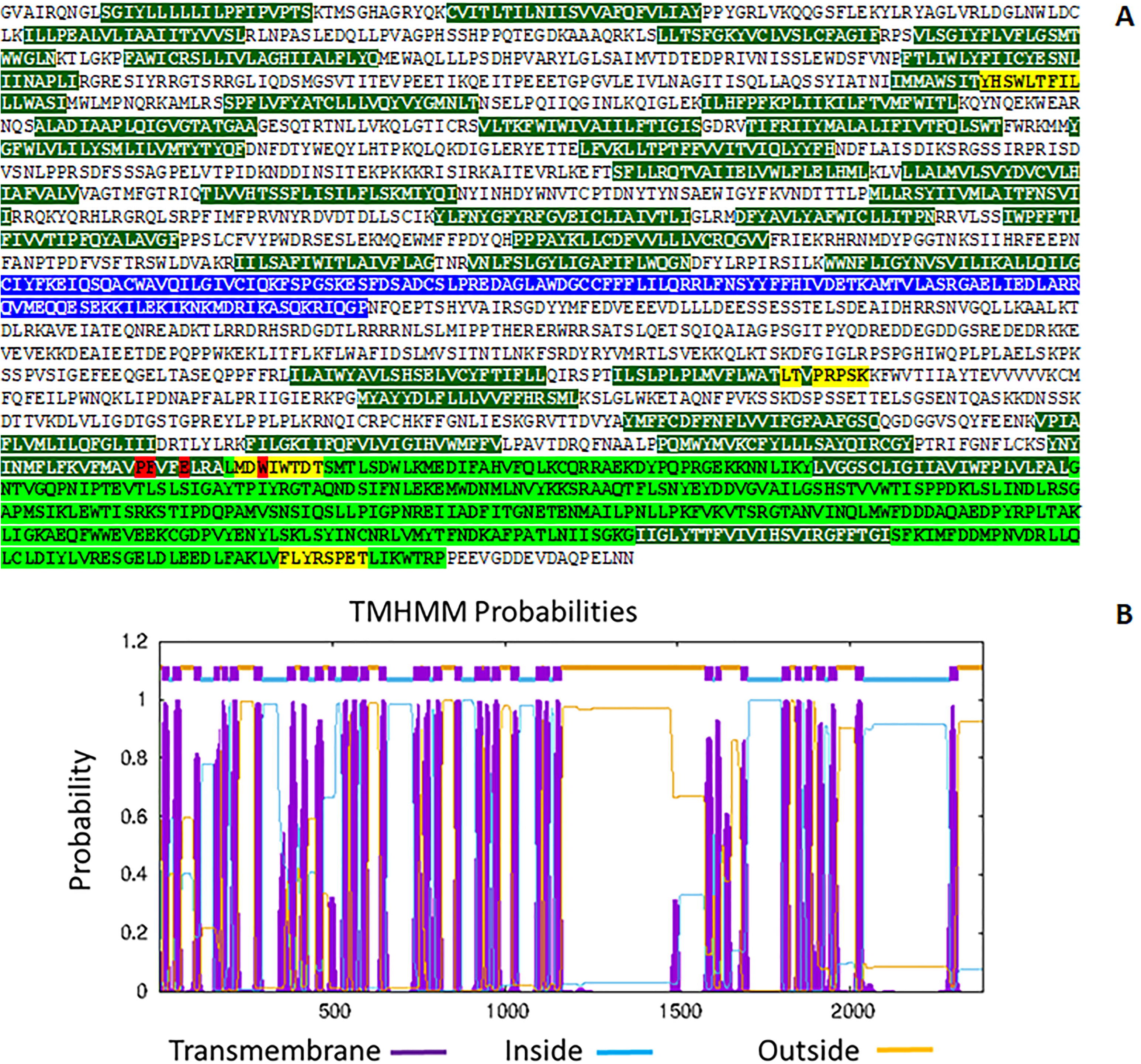
Piezo-like protein in *Rhodnius prolixus*. A) Sequence of a Piezo-like protein predicted by *R. prolixus* genome. Residues highlighted in blue and green represent conserved characteristic Piezo protein domains. Residues highlighted in yellow correspond to highly conserved motifs in Metazoa. Residues highlighted in dark green correspond to every transmembrane domain predicted in (B).

**Figure 8.**
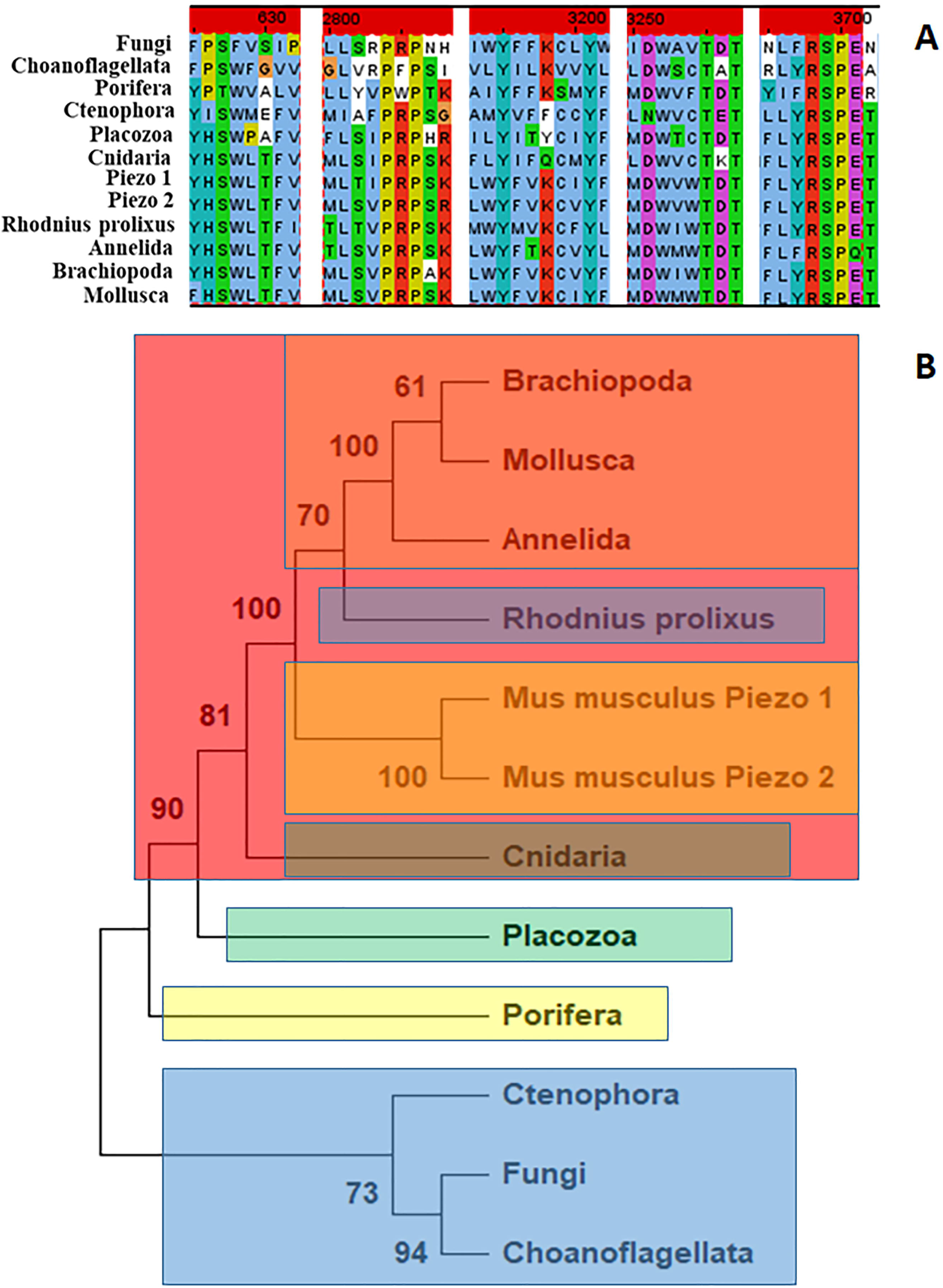
Homology and phylogenetic relationships between *R. prolixus* and several species pertaining to different phyla of Opisthokonta, including Metazoa, Cnidaria, Placozoa, Porifera and Ctenophora. Fungi and Choanoflagellata are also represented. The tree was constructed by Maximum Likelihood methodology (500 bootstraps).

The use of two different Piezo protein agonists (Jedi 1 and Jedi 2) physiologically confirm the presence of this family of protein as it is suggested by the increase of activity of both, the aorta and crop (Fig. 9 A and B). When the activity of the aorta was evaluated, results showed that both agonists (Jedi1 and Jedi2) caused a similar response. On the contrary, regarding the peristaltic activity of the crop, Jedi1 caused a major increase while Jedi2 induced a slight but still statistically significant response (Fig. 9 B).

**Figure 9.**
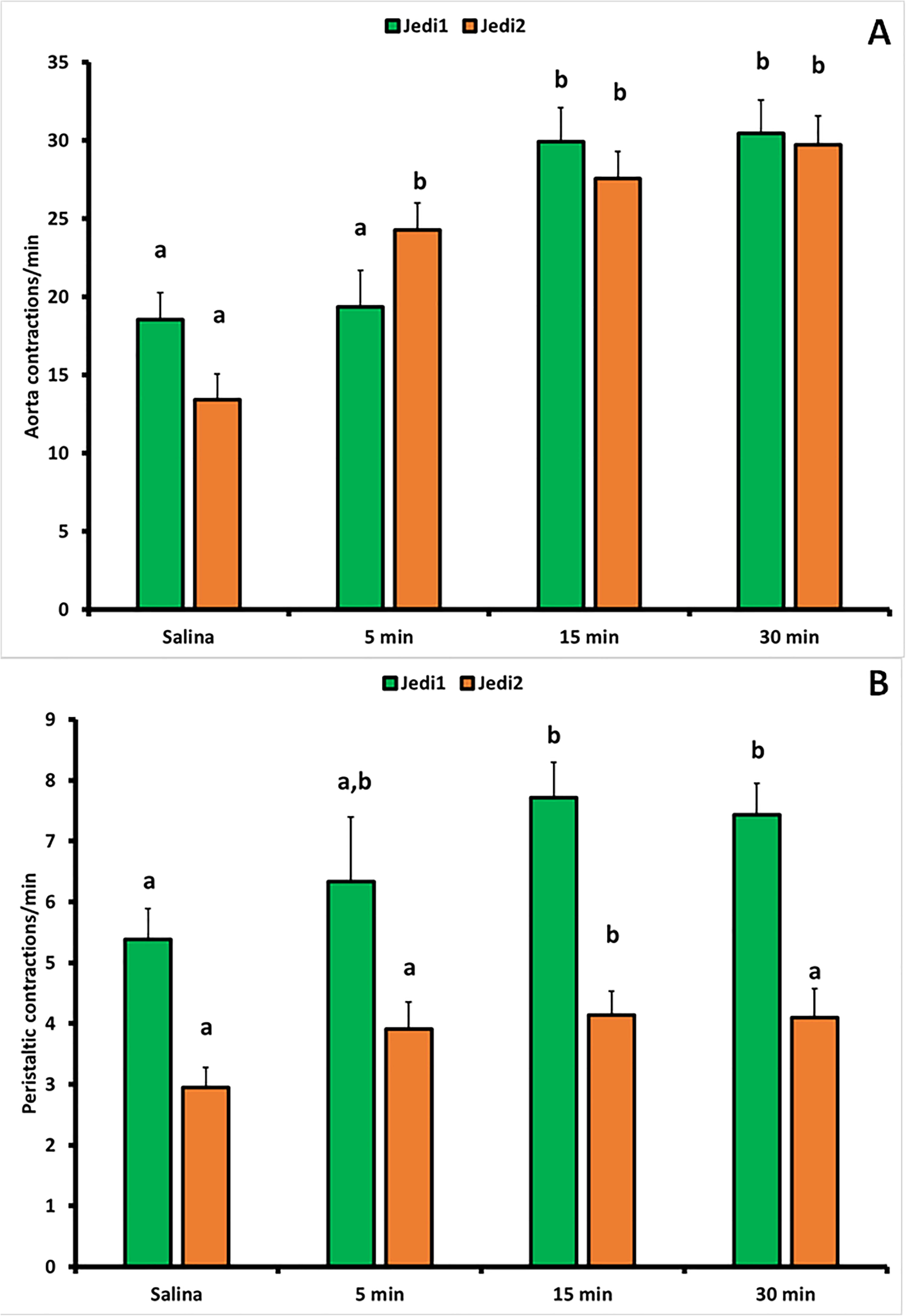
Effect of Jedi1 and Jedi2 Piezo proteins agonists in *R. prolixus*. A) Stimulatory activity of Piezo agonists on the frequency of contractions of the aorta. B) Effect of Jedi1 and Jedi2 on the peristaltic rate of contractions of the anterior midgut. Each bar represents mean ± SE. Different letters mean statistically significant differences.

## DISCUSSION

Triatominae insects ingest several times its own volume during feeding (Buxton, 1930). In order to concentrate the blood ingested and restore homeostasis, a physiological stress leading to an intense diuretic process during the next few hours occurs. Then, neuroendocrine mechanisms modulating moult in juvenile stages, and reproduction in adults begin. In fact, it was shown that the activity of the corpora allata in *T. infestans*, and the titre of JHSB3 in the hemolymph of *R. prolixus*, increase after feeding (Ronderos, 2009; Villalobos Sambucaro et al., 2020). To ensure diuretic activity, the increase of the velocity of circulation of the hemolymph must be guaranteed. Indeed, soon after the insect begins to feed, the rate of contractions of both, DV and crop, significantly increase (Maddrell, 1963, 1964a, 1964b). In triatominae insects, the physiological changes associated to diuresis (Maddrell, 1993a, 1993b) and those ones that modulate the frequency of contractions of the DV, and the rate of peristaltic activity of the crop have been extensively studied *in vivo* (Santini and Ronderos, 2007, 2009a; Sterkel et al., 2010; Villalobos Sambucaro et al., 2015, 2016). On the contrary, the information related to the mechanisms that modulate the hemolymph flow during the resting conditions are poorly understood. Triatominae insects can spend several weeks without feed, being processes such as diuresis, moult, copula and oviposition limited to short periods along its lifetime. In this regard it might be assumed that the resting situation would be the most frequently physiological condition along the life. Regarding that, the knowledge about the mechanisms ruling that phenomena, must be of interest to develop new approaches tending to the control of triatominae populations. In the present study we analysed several different pathways to delve into the mechanims ruling hemolymph circulation during the resting conditions in *R. prolixus*.

As we stated above, circulatory system in triatominae is simple, pumping the hemolymph along the abdomen and thorax, opening at the level of the head bathing the brain. The return is caused by peristaltic contractions of the anterior midgut. Regarding the myogenic contractions of the DV, there are not specific studies in *R. prolixus*, but the existence of heart automatism in insects has been proposed by several studies along the years (Maloeuf, 1935; Markou and Theophilidis, 2000; Sláma, 2006, 2012; Sláma and Lukás, 2011).

As a first attempt to understand the processes ruling the contractile activity of the aorta, we decided to check the relevance of Ca^2+^. It was evident that an increase of the cytosolic concentration is necessary as it was seen after chelate intracellular Ca^2+^ by mean of a permeable compound as BAPTA/AM. Moreover, as in other insect species (Markou and Theophilidis, 2000; Feliciano et al., 2011), our results demonstrated the Ca^2+^ influx is necessary to maintain the frequency of contractions. The relevance of the extracellular calcium influx was also confirmed by the use of a specific blocker of L-type VGCC (nifedipine) supporting the existence of action potentials driving the basal activity of the aorta. One of the most common response to the increase of cytosolic Ca^2+^ by mean of extracellular influx, is the further increase of the concentration of this ion due to its release from the ER through specific channels as RyR. When these channels were blockaded the contractile activity of the aorta also decreased. Indeed, when the DV is maintained *in vitro* a rhythm of contractions is still evident (Supplementary movie). All these results support the influence of a myogenic mechanism on the basal rhythm of contractions of the DV in *R. prolixus*.

Behind the myogenic nature of DV beating in insect, it is known that the rhythmicity is also modulated by neuropeptides and neurotransmitters (for a review see Hillyer, 2018). In *R. prolixus* and *T. infestans*, it was shown by in vivo experiments that, during post-prandial diuresis, 5-HT, AT and AST-C interact to regulate the speed of circulation of the hemolymph (Sterkel et al., 2010; Villalobos Sambucaro et al., 2015, 2016). To analyse the probable incidence of cellular messengers during the resting conditions, we assayed the effect of two compounds that alter a transductional pathway associated to GPCRs, such as Xe-C and U73122. Our results show that Xe-C, a natural compound that inhibits the opening of IP_3_R channels at the level of ER, causes a transient but statistically significant decrease of the frequency of contractions of the aorta, suggesting the involvement of a GPCR mediated signal. On the contrary U73122 which inhibits PLC (the enzyme responsible of the synthesis of IP_3_), did not cause a significant effect. Interestingly, a study showing a similar result on the spontaneous rhythmic contractions of the lateral oviduct of *Grillus bimaculatus* was previously reported (Tamashiro and Yoshino, 2014). To go further in the understanding of the peptides/GPCR pathway, we analysed the effect of an allosteric modulator of GPCRs. This compound inhibits ligand binding to GPCRs altering the activity of both stimulator and inhibitor ligands in vertebrates and invertebrates (Alzugaray et al., 2021, Fawzi et al., 2001; Gao et al., 2004; Hartz et al., 2008; Lewandowicz et al., 2006). Our results showed that insects treated with SCH-202676 underwent an increase of the frequency of contractions, suggesting the interaction between stimulatory and inhibitory signals. It was previously reported that AT and AST-C interact to modulate the frequency of contractions of the aorta during post-prandial diuresis, being the stimulatory effect of AT antagonized by AST-C (Villalobos Sambucaro et al., 2016). It was also reported that, *in vivo*, the myostimulatory activity of AT is mediated by the presence of 5-HT, having this peptide no effect when is applied alone (Masood and Orchard, 2014; Sterkel et al., 2010; Villalobos Sambucaro et al., 2015). Regarding this, we decided to check the probable existence of this functional relationship between AT and AST-C also under resting conditions. The results showed that in the presence of an anti-AST-C antiserum, AT was able to stimulate the frequency of contractions of the aorta, bringing an explanation for the effect caused by the GPCR allosteric modulator SCH-202676. This result also show that AT/AST-C interaction might be a canonical pathway acting to regulate and maintain the homeostasis of the visceral muscle activity at the level of the aorta and other organs under different physiological conditions. In fact, a similar mechanism was recently proposed in the female reproductive system (Villalobos Sambucaro et al., 2023). As stated above, it was previously shown that, under *in vivo* conditions, AT is not able to modulate contractile activity of the aorta when serotonin is not present (Masood and Orchard, 2014; Sterkel et al., 2010; Villalobos Sambucaro et al., 2015). Regarding the results obtained in this series of experiments, and taking into account that the aorta expresses both, AT and AST-C receptor (Villalobos Sambucaro et al., 2015, 2016), we assayed now the response of the aorta in the presence of AT (10^-9^ M) *ex vivo* (i.e. with no influence of the nervous and neuroendocrine systems) (supplementary movie 1). It is now clear that the aorta reacts to AT even without 5-HT, confirming that it is able to respond to the peptide, and that *in vivo* some kind of cross-talk between 5-HT and AST-C modulates its activity. AST-C activity is mediated by a GPCR homologue to somatostatin receptors in vertebrates (Alzugaray et al., 2016; Birgül et al., 1999). Indeed, a similar functional relationship between 5-HT and Somatostatin was proposed in vertebrates. It was shown that serotonin inhibits directly or indirectly somatostatin secretion both at the level of the stomach and pancreatic islets (Almaça et al., 2016; Koop and Arnold, 1984).

The frequency of contractions of the aorta is modulated by both myogenic as well as cell messenger signals. Neither the compounds that alter action potential response nor those associated to GPCR signalling, completely avoid the contractile activity by their selves. Regarding that, and trying to go further in the understanding of the mechanisms that control hemolymph circulation, we attempted to induce a complete blockage of the activity by treating the insects simultaneously with those compounds that proved to alter the contractions (i.e. Ry, Xe-C and nifedipine). Again, a complete blockage of the contractile activity was not evident. These results suggest that behind automatism and neuropeptides and neurotransmitters signalling, another via should be involved to ensure at least a minimal function of the circulatory circuit.

As in previous studies, we also evaluated the peristaltic rate of contractions of the anterior midgut. As expected, due to the relevance of calcium during muscle contraction, the system resulted to be sensitive to the lack of this ion. This was evident after treatment with both extra and intracellular chelators. On the contrary, only Ry was effective altering peristaltic activity. Interestingly, the opening of RyRs did not show to be associated to L-type VGCC, as nifedipine did not cause any effect. Taking together, these results suggest that other membrane associated proteins might be involved facilitating calcium income to open RyR and to induce muscle contraction. Moreover, not myogenic activity seems to be present on the crop. Furthermore, contrarily to the aorta, crop behaviour seems to be variable and aleatory. As the DV pumps regularly in an anterograde direction causing an increase of the hemolymph at the level of the thorax and the head, this increment could cause a slight but significant increase of the hemolymph pressure to trigger a reflex contraction of the muscle wall. Indeed, it was demonstrated in the related species *T. infestans*, the existence of allatotropic open type cells in the anterior region of the crop acting in a paracrine way (Riccillo and Ronderos, 2010; Sterkel et al., 2010). The pressure caused by the hemolymph accumulated could stimulate the release of AT triggering a peristaltic wave. To delve into the mechanisms and to explore this possibility, we look for the existence of Piezo proteins in the genome of *R. prolixus*. Piezo proteins are a recently characterized family of proteins associated to cell membrane. They are sensitive to mechanical forces, facilitating the influx of cations (Coste et al., 2010, 2012) including Ca^2+^ (Zechini et al., 2022) and they were associated to multiple functions (see Wu et al., 2017). Originally characterized in vertebrates, they were also found in invertebrates. In *D. melanogaster*, these proteins proved to be involved, in nociception (Kim et al., 2012), control of meal size (Min et al., 2021) and cardiomyocytes activity (Zechini et al., 2022).

Looking for a probable reflex response mediated by this way, we searched for Piezo-like proteins in the genome of *R. prolixus*. We found that *R. prolixus* genome predicts the expression of a protein sharing a high level of homology with vertebrates. Indeed, Piezo proteins are a highly conserved family being present in most eukaryotes including plants and protists (Coste et al., 2010).

Regarding *R. prolixus*, we found a protein which share two recognizable domains including the one associated to de C-terminal, which in fact contains pore properties (Coste et al., 2015). We found several other motifs highly conserved such as FLYRSPET which in fact could be considered as a signature for Metazoa. The use of two agonists as Jedi 1 and Jedi 2, induced a statistically significant increase of the frequency of contractions at both, aorta and anterior midgut, suggesting the expression of Piezo proteins in vivo. Indeed, due that the specificity of action of these agonists, it should assumed that the Piezo protein present in *R. prolixus* should be a Piezo 1 protein (Wang et al., 2018).

Our results show that under resting conditions, the mechanisms controlling hemolymph circulation, including the ones acting on the anterograde flow (i.e. DV pumping) and the retrograde movement (peristaltic activity of the crop) are highly complex. Indeed, they include several different factors including automatism, nervous, neuroendocrine and also paracrine pathways. Certainly, behind of those ones, and as a mechanism to ensure the maintenance of a minimum flow in resting conditions, the relevance of a reflex mechanism drive by a mechano-sensitive system as Piezo proteins is present.

## ACKNOWLEDGEMENTS

This work was supported by funds provided by the Universidad Nacional de La Plata (N/948) and FONCyT (PICT: BID-PICT 2018 N° 1236). MJVS and MEA are researchers at CONICET (Argentina). The authors wish to thanks to Dr. F.G. Noriega for generously supplying Allatotropin and Allatostatin-C.

## CONTRIBUTORS

Conceived and designed the experiments: JRR, MJVS; Performed the experiments: MJVS; Analysed the data: JRR, MJVS, MEA. Wrote the paper: JRR. Contributed reagents/materials/analysis tools: JRR, MJVS.

**Supplementary movie.** Allatotropin stimulates the contractile activity of the aorta *ex vivo*.

